# Top-down attention shifts behavioral and neural event boundaries in narratives with overlapping event scripts

**DOI:** 10.1101/2023.08.08.552465

**Authors:** Alexandra De Soares, Tony Kim, Franck Mugisho, Elen Zhu, Allison Lin, Chen Zheng, Christopher Baldassano

## Abstract

Understanding and remembering the complex experiences of everyday life relies critically on prior schematic knowledge about how events in our world unfold over time. How does the brain construct event representations from a library of schematic scripts, and how does activating a specific script impact the way that events are segmented in time? We developed a novel set of 16 audio narratives, each of which combines one of four location-relevant event scripts (restaurant, airport, grocery store, lecture hall) with one of four socially-relevant event scripts (breakup, proposal, business deal, meet cute), and presented them to participants in an fMRI study and a separate online study. Responses in angular gyrus, parahippocampal gyrus, and subregions of medial prefrontal cortex (mPFC) were driven by both location and social script information, showing that these regions can track schematic sequences from multiple domains. For some stories participants were primed to attend to one of the two scripts, by training them to listen for and remember specific script-relevant episodic details. Activating a location-related event script shifted the timing of subjective event boundaries to align with script-relevant changes in the narratives, and this behavioral shift was mirrored in the timing of neural responses, with mPFC event boundaries (identified using a Hidden Markov Model) aligning to location-relevant rather than socially-relevant boundaries when participants were location primed. Our findings demonstrate that neural event dynamics are actively modulated by top-down goals, and provide new insight into how narrative event representations are constructed through the activation of temporally-structured prior knowledge.

## Introduction

A key challenge in human neuroscience is to understand how the complex, continuous stream of information we experience during our everyday lives is used to construct stable and meaningful neural representations. Work over the past decade has shown that, for stimuli with meaningful temporal structure (such as narratives or film clips), rapidly-changing responses in early sensory regions are transformed into slower, more stable, and more abstract representations through a series of regions along a temporal processing hierarchy ^1–4^. At the top of this hierarchy, in regions traditionally characterized as the Default Mode Network (DMN), neural representations exhibit periods of relative stability (for tens of seconds) punctuated by moments of rapid change ^5, 6^. These dynamics align with a long-standing idea in cognitive psychology: that continuous experiences are segmented into discrete, meaningful events ^7–10^, with boundaries between events serving as critical moments for updating mental models in working memory ^11–13^ and for determining the structure of episodic memory ^14–17^.

Both the content of event representations and the timing of event boundaries are thought to be scaffolded by the activation of schematic event scripts ^18–20^ that reflect prior knowledge about the sequential temporal structure of events we encounter in the world, allowing us to strategically direct our attention and make predictions ^21–23^. For example, when catching a flight, experienced travelers will have clear expectations about the sequence of events that will occur at an airport and the critical pieces of information they will need to track at each stage. Previous work using naturalistic events has specifically implicated the medial prefrontal cortex (mPFC) in tracking this abstract event structure: mPFC responds more strongly to schema-consistent events ^24^ and has representations that generalize across narratives with shared event schemas ^25–28^, and mPFC lesions impair the activation of schematic event knowledge ^29^.

Prior neuroimaging studies of event scripts, however, have had two major limitations. First, these studies have assumed that there is only a single relevant script active during a story, while realistic events are much more likely to be a combination of multiple, overlapping scripts, such as celebrating a birthday at a restaurant. Even the earliest work on story scripts noted that “the concurrent activation of more than one script creates rather complex problems” ^18^; multiple scripts must be simultaneously tracked, scripts can compete for incoming pieces of information and for attention, and different scripts may generate conflicting event boundaries. One possibility for how the brain could track multiple scripts simultaneously is by representing different kinds of scripts in different brain regions, consistent with prior work showing that specific subnetworks of the DMN are most engaged by particular kinds of tasks, such as tracking mental states or spatial imagery ^30–32^. Alternatively, knowledge from multiple scripts (along with episode-unique details) could be integrated into a single high-dimensional event model in DMN regions ^33–35^, especially mPFC ^25, 36, 37^.

The other major shortcoming of previous paradigms for studying event scripts is that they cannot address the central theoretical claim that top-down script activation impacts how neural responses are organized into events ^15, 23, 38^. A script template could be used to actively stabilize an event representation over time, by ignoring sensory changes irrelevant to the current event type ^15^; an alternative explanation, however, is that the observed event dynamics are already present in the stimulus itself, and that simply detecting temporally-stable features in the stimulus (such as locations) would yield stable patterns of neural activity ^39^. Narratives with multiple simultaneous scripts provide an intriguing opportunity to examine the causal impact of activating an event script on neural responses, since the level of attention to each script can be manipulated while holding the narrative stimulus constant. Previous work has shown that activating a (static) schema during a narrative impacts perception and memory, improving memory for details relevant to the primed schema ^40^ and also modulating responses in the DMN^41–43^. Here we tested whether priming participants to sequentially attend to each stage of an event script can cause a corresponding change in the timing of neural event boundaries. We hypothesized that encountering a narrative event boundary relevant to the currently-active script will generate an internal state shift analogous to switching between task sets, which has been shown to generate event boundaries in memory ^44^.

Our study aimed to address these questions about how multiple scripts are tracked during narrative perception and how directing attention to a specific script can influence perceptual dynamics. We collected fMRI data while participants listened to a novel set of narrative stimuli in which each story was built from two distinct event scripts of four schematic events each (Figure 1). One of these scripts was determined by the *location* that the story took place (e.g. an airport), and the other described the *social* interaction taking place (e.g. a marriage proposal). There were four location scripts and four social scripts, and each of the sixteen stories had a unique combination of a location and social script, interleaved such that the event boundaries for each script occurred at different sentences during the story. We found that regions throughout the DMN integrated information about both location and social scripts. We also manipulated attention during some of the stories (in our fMRI study and in an online study) by asking participants to detect and remember a sequence of details relevant to one of the scripts, priming them to consecutively activate the events of either the location script or social script. We found that activating the location-relevant script shifted subjective event boundaries to align with location-relevant boundaries in the narrative, and that there was a corresponding shift in the timing of neurally-identified event boundaries in mPFC. These results provide a key link connecting neural dynamics during naturalistic perception to decades of work on schematic perception and event segmentation in cognitive psychology, demonstrating that event boundaries are actively constructed and not simply produced by stimulus dynamics.

**Figure 1:**
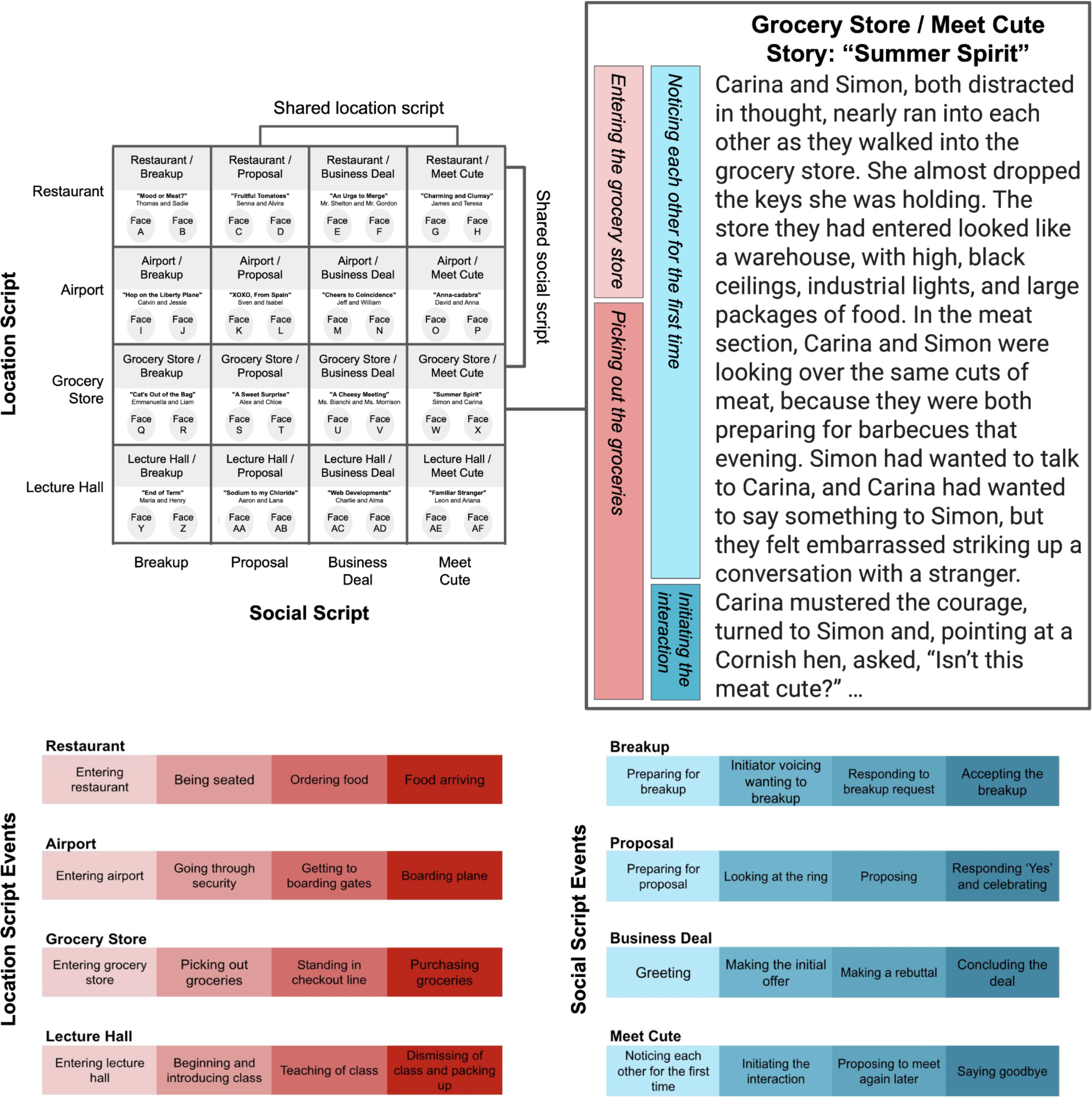
Narrative stimulus set with overlapping scripts. We selected four location-based scripts and four social-interaction scripts, each of which consisted of a stereotyped four-stage sequence of events. Each story was written based on a unique combination of a location and social script, resulting in four narratives for each script. The events in both scripts occurred simultaneously during the narrative, with event boundaries interleaved such that script-relevant transitions occurred at different sentences for the two scripts. The story was presented as spoken audio, while a static image of the story title and the two characters’ faces and names was displayed. See Supplemental Figure S1 for an illustration of event timings for all stories.

## Results

### Location and social scripts are represented in overlapping regions

To identify brain regions (in searchlights and predefined ROIs) that represented information about the location and social scripts, we measured the across-participant similarity for event patterns evoked by stories with shared scripts (Figure 2A). Both location-related and social-related scripts were represented in overlapping regions throughout the DMN, particularly in the middle temporal cortex (MTC), the right angular gyrus, parahippocampal cortex (PHC), and a dorsal portion of mPFC (Figure 2B). We also saw script-specific event representations for both script types in inferior frontal gyrus and in parts of visual cortex, despite there being no dynamic visual content in these stimuli. There were some additional regions that represented only a single category of scripts, with selectivity for location-script effects in a broader set of visual regions and left insula and social-script effects in the left temporoparietal junction, left posterior medial cortex (PMC), and broader mPFC. Measuring the location and social script effects in predefined ROIs (Figure 2C), we found overlapping representations in the angular gyrus (location script effect: p = 0.008; social script effect: p < 0.001) and PHC (location p = 0.004; social p = 0.012). Significant social script effects (but not significant location script effects) were observed in the MTC (location p = 0.17; social p = 0.013), superior frontal gyrus (SFG; location p = 0.51; social p = 0.002), mPFC (location p = 0.128; social p = 0.018), and hippocampus (location p = 0.153; social p = 0.010). We also conducted additional analyses of univariate effects in the hippocampus, finding increases in activity in response to the critical episodic details in each narrative as well as after event boundaries (see Supplemental Figure S2). Overall, our results show that there is some variability in the DMN regions that track location and social scripts, but that key regions including angular gyrus, PHC, and dorsal mPFC build representations that draw on both kinds of scripts in a domain-general way.

**Figure 2:**
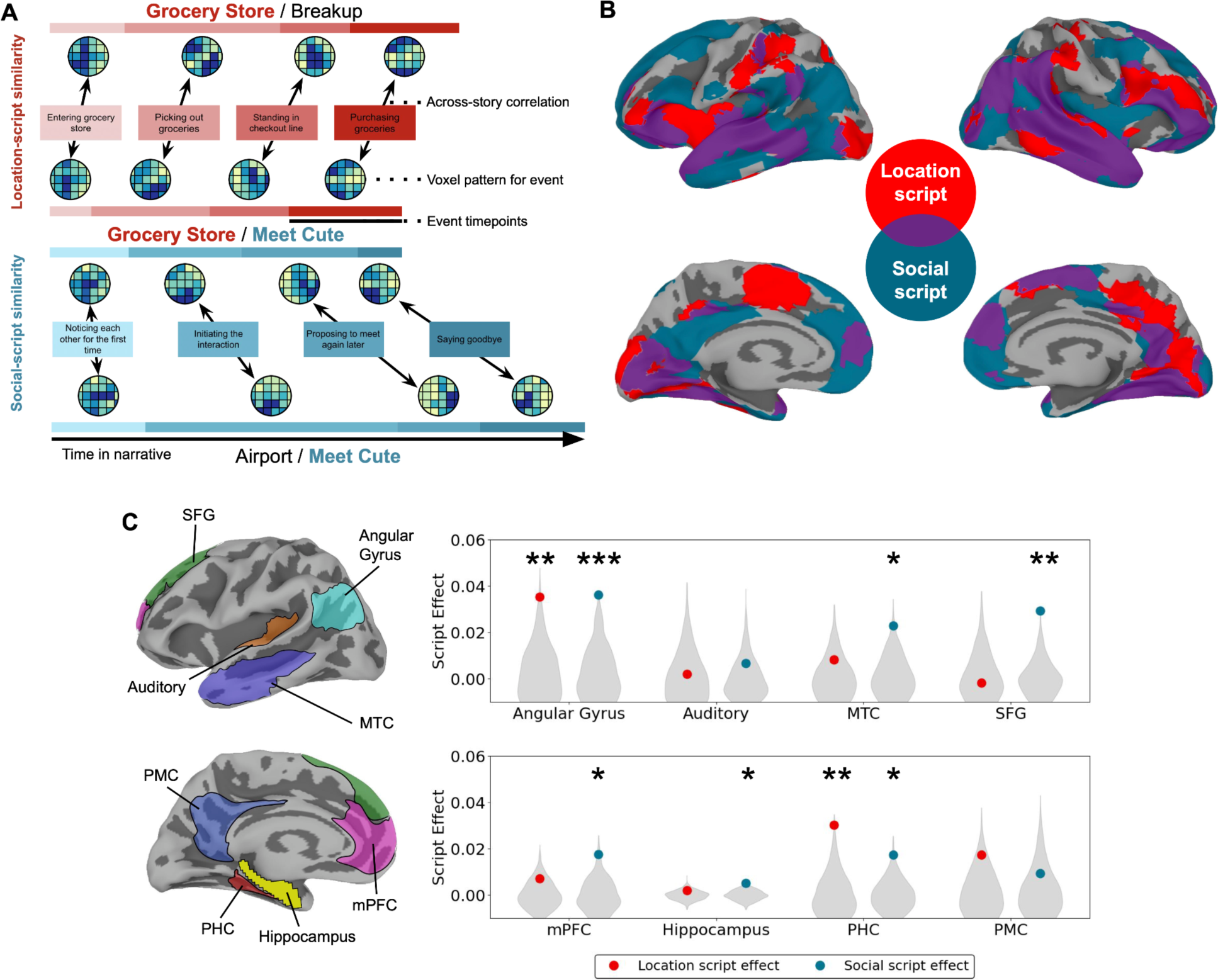
Location and social script effects in searchlights and ROIs. (A) We compared event patterns across stories with shared scripts to identify regions that encode script-specific information. For each story (e.g. the Grocery Store / Meet Cute story), we computed the average voxel activity pattern in a region for each location-script (e.g. Grocery Store) event and then compared corresponding event patterns across stories with shared location scripts (e.g. other Grocery Store stories). We then repeated the analysis, defining event patterns based on social-script (e.g. Meet Cute) events and testing for similarity across stories with shared social scripts (e.g. other Meet Cute stories). (B) Performing a searchlight analysis on the cortical surface, we identified a broad network of regions (purple) in which representations were influenced by both kinds of scripts, including bilateral portions of mPFC, PHC, inferior frontal gyrus, and lateral temporal cortex. There were also some regions (red and blue) that tracked only one category of scripts. Both maps are thresholded at q < 0.05. (C) Computing script effects in predefined ROIs ^25^, we found that angular gyrus and PHC significantly encoded both kinds of scripts, while the MTC, SFG, mPFC, and hippocampus significantly encoded the social script only. See Supplemental Figure S2 for additional analyses of hippocampal responses. * p < 0.05, ** p < 0.01, *** p < 0.001

### Biasing attention to an event script impacts memory and segmentation

To what extent does the activation of scripts depend on top-down attentional mechanisms, and what are the behavioral consequences of script activation? We conducted a separate online behavioral study (N=308) in which we primed participants to attend to either the location or social script in one of our sixteen stories (or a baseline condition, in which they did not receive either prime). We asked them to take on the perspective of a role associated with the schema (e.g. restaurant critic role for the restaurant script), and directed attention to each of the four script events by providing a sequence of four questions they should aim to answer while reading the story (Figure 3). The choice of which story was presented and the assignment to one of the three priming conditions was randomized across participants.

**Figure 3:**
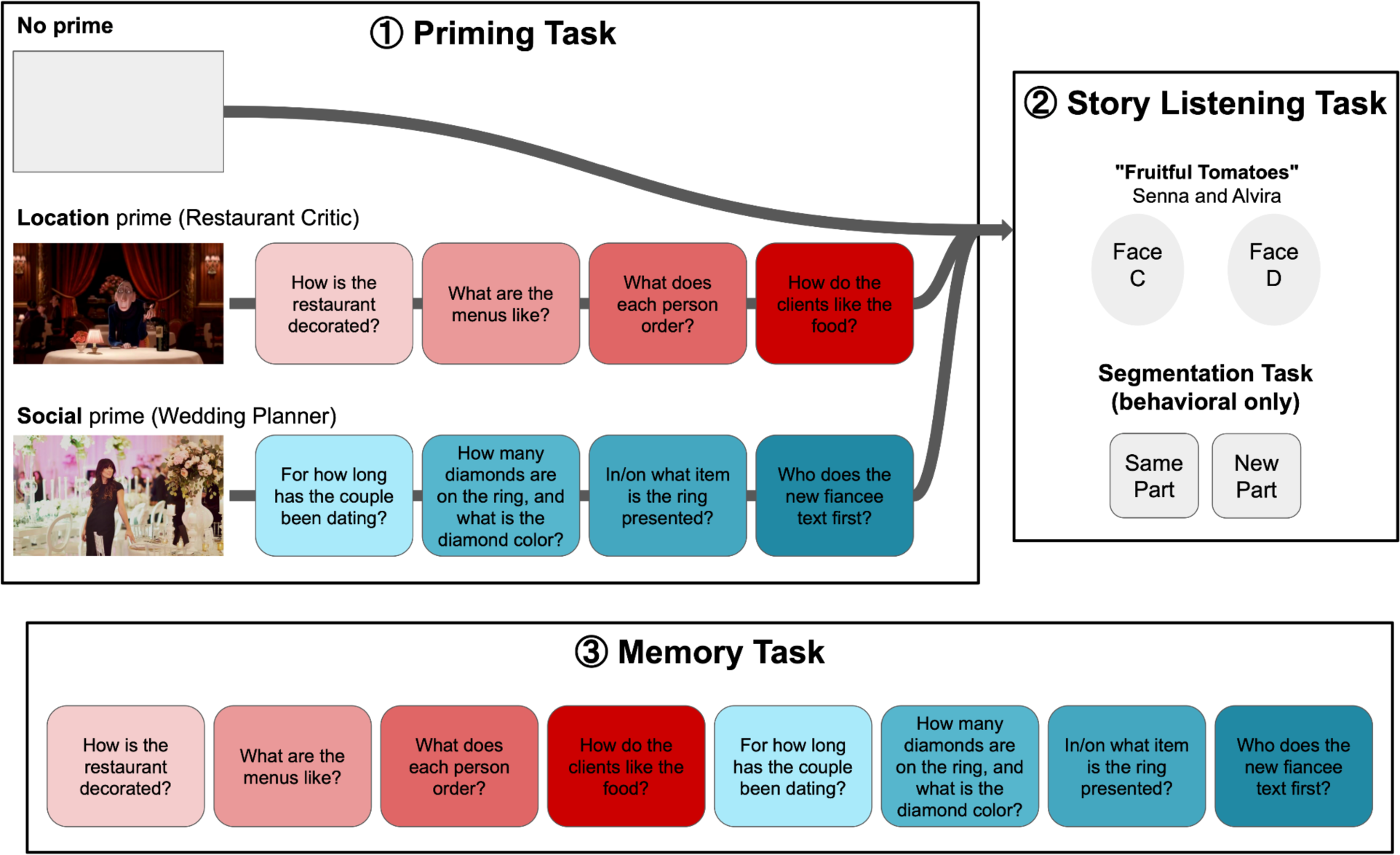
Event script priming and behavioral segmentation task. In both our fMRI experiment and a separate behavioral experiment, we placed participants in one of three priming conditions: no prime, location prime, or social prime. Participants in the location or social prime condition were told to take on the perspective of a role associated with the corresponding event script; examples are shown for one location-relevant role (restaurant critic) and one socially-relevant role (wedding planning). We specifically directed their attention to each event in the primed script by providing them with a sequence of 4 questions they should aim to answer while listening to the story. Participants had to correctly identify and order the questions in the priming task in order to move on to the Story Listening Task, which was identical for all priming groups. In the online study only, participants completed a Segmentation Task while they listened to the story, indicating after listening to each sentence if that sentence was a “new part” of the story. Participants were never exposed to the perspective questions for the unprimed scripts in their stories; online participants heard only one story, and fMRI participants heard only 12 stories in which they were primed for at most one of the scripts. At the end of the experiment, participants answered eight questions about story-specific details for each story they heard, with four questions related to each script.

Using a memory test at the end of the story, we confirmed that script priming impacted the episodic details that participants were able to recall. Participants were asked questions about episodic details in the story they had just heard and were scored on a scale of 0-3 for how precisely they could recall the correct answer (Figure 4A). A mixed-effects model showed that social-script priming improved performance specifically on socially-relevant questions (β = 0.393, t_372.00_ = 4.096, p < 0.001), and there was also better overall performance for location-primed participants (β = 0.262, t_628.29_ = 2.964, p = 0.003) and on socially-relevant questions (β = 0.266, t_372.00_ = 4.107, p < 0.001). Directly comparing only the location- vs. socially-primed participants with an independent-samples t-test, we found superior performance on details relevant to the location script for location-primed participants (Δμ = 0.234, t_231.34_ = 2.620, p = 0.009) and on details relevant to the social script for social-primed participants (Δμ = 0.277, t_227.76_ = 2.620, p = 0.007). We replicated these memory results in our fMRI participants (Figure 4B), finding improved scores on socially-relevant details with social priming (β = 0.638, t_823_ = 5.700, p < 0.001) and improved performance on location-relevant details for location-primed participants (β = 0.461, t_823_ = 4.122, p < 0.001), with better overall performance for location-primed participants (β = 0.559, t_823_ = 7.066, p < 0.001) and better scores for socially-relevant questions (β = 0.343, t_823_ = 4.338, p < 0.001). Directly comparing the location and social priming conditions only, we found significantly better performance on location questions for location-primed participants (Δμ = 0.517, t_251_ = 6.686, p < 0.001) and on social questions for socially-primed participants (Δμ = 0.582, t_251_ = 6.767, p < 0.001).

**Figure 4:**
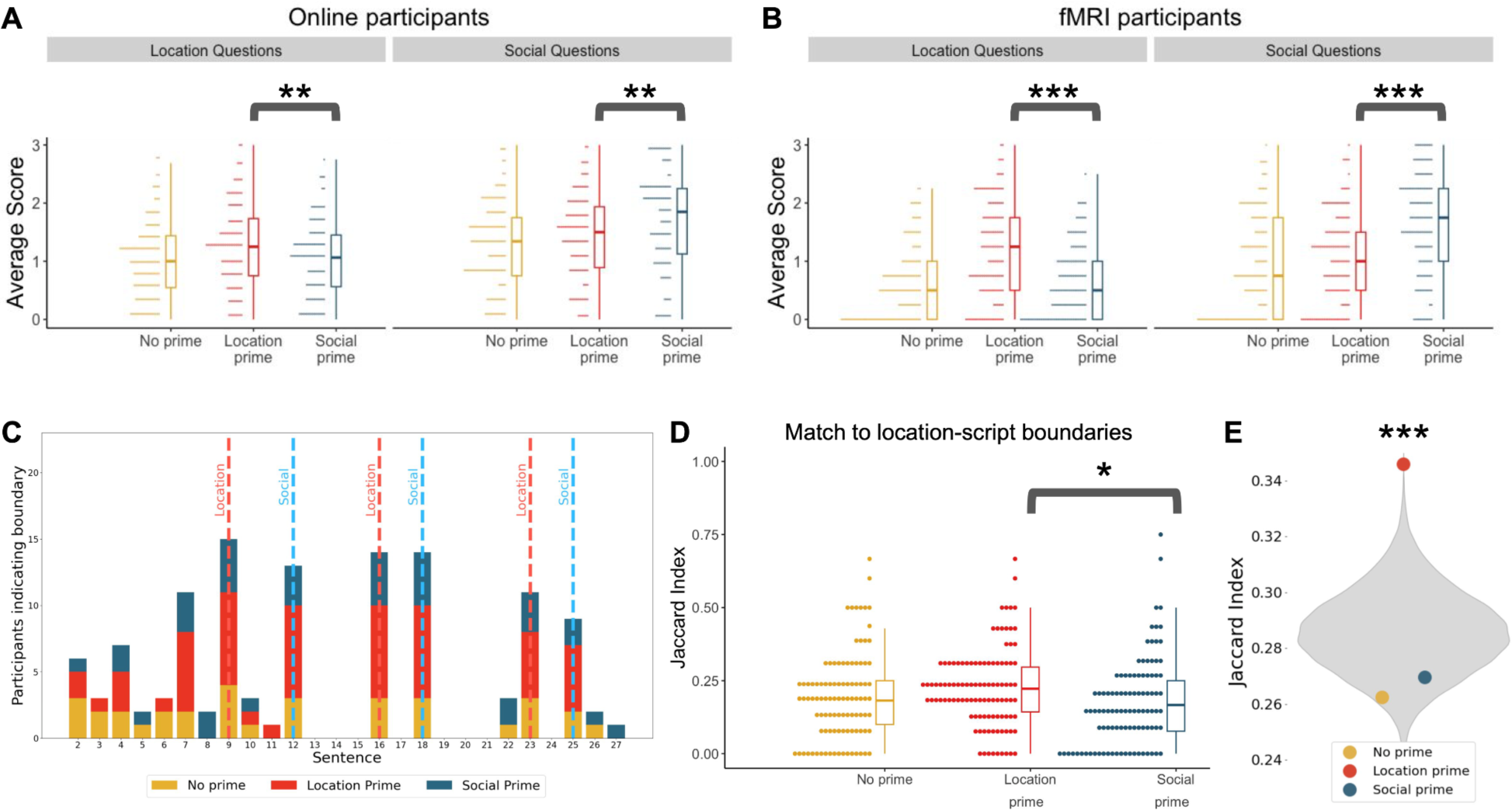
Behavioral effects of biasing attention to event scripts. (A) Online participants were scored on their recall of story-specific details relevant to the location and social scripts. Priming had a significant effect on how participants remembered story details, with better memory for details relevant to the primed script. (B) This effect replicated in our fMRI participants, who also showed significantly better memory for prime-relevant details. (C) Online participants in all priming groups showed substantial agreement about which sentences were event boundaries, shown here for one example story. Colored bars indicate the number of participants in each condition that labeled a sentence as starting a new event, with dotted lines denoting the event transitions for each script. (D) Boundaries annotated by location-primed participants were significantly more aligned to the location-related transitions in the narrative, compared to social-primed participants. (E) Location-primed participants were significantly more similar to each other compared to random pairs of participants (gray). See Supplemental Table S3 for additional analyses of boundary responses. * p < 0.05, ** p < 0.01, *** p < 0.001

We also asked online participants to identify event boundaries while reading the story, indicating whether each sentence was the start of a new event (Figure 4C). Participants in all priming groups showed substantial agreement about which sentences were event boundaries, relative to a null distribution of boundaries with a matched distribution of event lengths (p < 0.001 for all groups). All priming conditions yielded similar numbers of event boundaries (no prime: 6.79; location prime: 7.64; social prime: 7.03; ANOVA for effect of priming condition, F_2,305_=1.71, p=0.183). To determine whether the priming caused a systematic shift in boundary judgments toward the script-related boundaries we had created in the stories, we compared participants’ event boundaries to the putative event boundaries corresponding to each script (Figure 4D). We found that the priming groups differed in their alignment to the location boundaries (F_2,305_ = 3.53, p = 0.030), with significantly better alignment for participants who were location-primed rather than socially-primed (p = 0.023 by post-hoc Tukey test). We did not observe differences across priming groups for alignment to the social-script boundaries (F_2,305_ = 0.26, p = 0.77). We also tested whether the event boundaries of participants who were primed in the same way became more similar to each other (regardless of whether they matched our putative script-related boundaries in the stimulus). We found (Figure 4E) that location-primed participants had boundaries that were significantly more similar than for random pairs of participants (p < 0.001), while social-script priming did not significantly improve boundary similarity (p = 0.860). These results show that manipulating script attention can influence the episodic details remembered in a narrative, and can (for location-related scripts) align subjective event boundaries to script-relevant event changes.

### Priming shifts the timing of event boundaries in mPFC

Since our online study showed that biasing attention toward a location-related script shifted subjective event boundaries to align with location-script stimulus boundaries, we tested whether there was a corresponding shift in neural event boundaries. We fit a four-state Hidden Markov Model (HMM) to the story-evoked activation for each priming group, temporally clustering multivoxel responses into stable events separated by rapid transitions (Figure 5A). The HMM therefore allowed us to derive a measure of neural event change at each timepoint, for each of the priming groups (Figure 5B). We measured the average neural event change at the putative location and social event boundaries to produce a measure of alignment between neural event boundaries and putative event boundaries (for an alternative measure of neural change at event boundaries, see Supplemental Figure S3).

**Figure 5:**
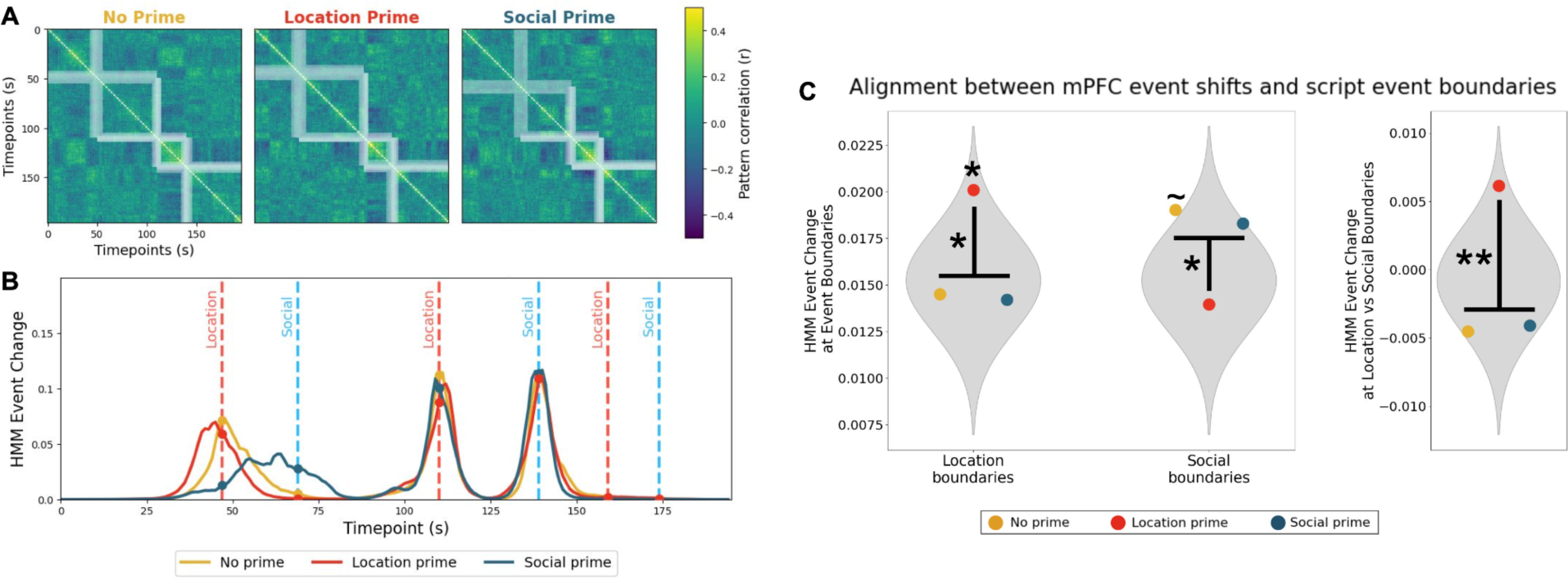
Alignment between mPFC events and script event boundaries across priming groups. (A) The matrix for each priming group shows the correlation between mPFC voxel patterns for all pairs of timepoints in an example story. Blocks of high values along the diagonal indicate periods when activity patterns are remaining relatively stable over time. The HMM event segmentation model identifies moments of rapid event change (shown in white) between these stable events. (B) We can measure the alignment between event change in the brain and the location-script and social-script boundaries by computing the HMM event change at each boundary (dots). In this example story, we see that all groups have neural shifts that are well-aligned with boundaries in the story, but that social priming shifts the first neural boundary from a location-script boundary to a social-script boundary. (C) Across all stories, we find that location-primed participants are significantly aligned to location boundaries, and that location priming causes a significant increase in alignment to location boundaries and a significant decrease in alignment to social boundaries. Combining data from both kinds of boundaries, we find that location priming drives a significant overall effect in shifting alignment from social boundaries to location boundaries. See Supplemental Figure S3 for an alternative measure of neural change, Supplemental Figure S4 for results from additional ROIs, Supplemental Figure S5 for an alternative analysis of neural changes with priming, and Supplemental Figure S6 for analysis of sentences with episodic details. ∼ p < 0.10, * p < 0.05, ** p < 0.01

Our primary ROI of interest for this analysis was mPFC, based on extensive prior work implicating the mPFC in schema activation ^24–29, 45, 46^ (Figure 5C). Computing an overall measure of priming-driven boundary shifts (measuring the difference in alignment to location vs social boundaries when participants were location-primed, compared no priming or social priming), we found that neural event alignment shifted significantly from social boundaries to location boundaries after location-script priming (p=0.007). We also conducted exploratory analyses for each type of boundary separately. For location-script boundaries, only location-primed participants were significantly aligned (p=0.039; no-prime p=0.606, social-prime p=0.645), and this alignment was significantly greater than for other priming groups (p=0.013). For social-script boundaries, we observed the opposite effect, with significantly lower alignment for location priming (p=0.019) and marginal boundary alignment for no-prime participants (p=0.084; location-prime p=0.680, social-prime p=0.133). Because our prior analysis (Figure 2C) revealed that angular gyrus and PHC were sensitive to both location and social scripts (and therefore might be able to flexibly track either location or script boundaries in different priming conditions), we also ran this same analysis in those ROIs, and found an overall priming effect in PHC and differences in social-boundary alignment in both regions (see Supplemental Figure S4). These shifts were consistent with our behavioral results, showing that location script priming has an effect on the timing of neural event boundaries and that these boundaries become more aligned with event transitions relevant to the location-related script.

## Discussion

Using narratives that combined multiple kinds of event scripts, we found that scripts related to both physical location and social interactions influenced neural event representations of core event-perception regions in the Default Mode Network (DMN). Additionally, we used multi-script narratives as a tool to understand the top-down impact of script activation on perception and memory. By biasing attention to one of the scripts in the narrative, we showed that script activation changes behavioral outcomes, influencing subjective judgements of event boundaries and changing which episodic details are remembered. We also observed impacts of script activation in the brain, including event boundaries shifting to prime-relevant timepoints in regions including mPFC. These results argue for a view of event representation and segmentation as an active process, in which schematic event knowledge plays a key role in the dynamics of mental and neural representations during narrative perception.

### The Default Mode Network flexibly represents narrative scripts

While we found that script information was broadly distributed throughout the cortex, representations of *both* location and social scripts were focused specifically on regions in the DMN, especially angular gyrus, PHC, and portions of MTC and mPFC. Our study extends previous work that used a more limited set of event schemas (restaurant/airport ^25^ or cafe/grocery ^26^), demonstrating that schemas need not be tied to particular locations to be tracked by the DMN. These findings support the theory that DMN regions are not constrained to processing or representing particular semantic categories of information, but instead carry out domain-general abstract cognitive processes. In particular, angular gyrus has been proposed as a general-purpose buffer for maintaining event information over time ^47^ and in the PHC has been shown to perform contextual associative processing in both spatial and nonspatial tasks ^48^. Using familiar past experiences as building blocks ^49^ that can be flexibly combined and remixed across domains allows us to rapidly comprehend complex events, even when the information in the stimulus is highly restricted ^43^. The broader set of event scripts used in this study are of course only a small fraction of the full set of schemas employed in narratives; future work could examine a broader basis set of event schemas and account for idiosyncratic knowledge and expertise of specific participants.

More generally, our results support an emerging model of the DMN as a convergence zone integrating stimulus-driven signals with idiosyncratic internal knowledge about the world such as memories, schemas, and goals ^34^. Originally labeled as a “task-negative” network ^50^, the DMN is still often characterized as being engaged only for internally-driven cognition ^51^ or “internal mentation tasks” ^52^. This traditional view has been challenged by paradigms using more naturalistic stimuli that have long-timescale dependencies over tens of seconds and connect to real-world prior knowledge, which in fact robustly engage the DMN ^53–55^. We argue that the DMN is therefore better described as carrying out cognitive processes with meaningful semantic and temporal structure, regardless of whether these are externally or internally driven. The DMN representation of external narrative stimuli does differ from representations in sensory regions in its degree of sensitivity to prior experiences and prior context; providing alternative framings for a story can impact DMN responses throughout the entire narrative ^42, 56^, as we observed with our priming manipulation.

### Script representation in the hippocampus

We found that the hippocampus exhibited limited sensitivity to scripts compared to cortical DMN regions, with no significant tracking of location scripts and small but significant correlations for social scripts. These results are broadly consistent with past studies examining narrative schema effects, which have failed to find location-related script patterns in the hippocampus ^25^ or have found schema effects that were significantly weaker than episode-specific effects ^26^. These limited findings could stem from difficulties in measuring hippocampal activity with fMRI, due to acquisition difficulties in the medial temporal lobe ^57^ or highly sparse representations ^58^. An alternative is well-learned schemas, like the real-world scripts in our study, are in fact represented primarily by the cortex, with prefrontal regions (rather than the hippocampus) integrating information across cortical modules and possibly even inhibiting hippocampal responses to schematic elements ^59, 60^. For temporal scripts, however, this view appears at odds with findings that the hippocampus represents even well-learned temporal sequences ^61–63^.

We did observe some representation of social scripts in the hippocampus, suggesting that the hippocampus plays at least a partial role in tracking and/or encoding information about ongoing social interactions. While the hippocampus has traditionally been thought to be primarily oriented toward representations of physical space ^64^, it has also been hypothesized to represent more abstract structures like social information ^65^. The most general version of this hypothesis is that the computations in the hippocampus could support any kind of experience with sequential relational parts, compressing such instances to represent their latent structure ^66^. Future work on narrative scripts (perhaps using direct intracranial measurements of hippocampal responses to narratives in humans ^67^) could further investigate the role of the hippocampus in tracking abstract event types and how this role depends on task, consolidation over time, and expertise.

In our univariate analyses of hippocampal activity, we replicated past findings of increased activity following event boundaries ^5, 68–70^ and found that these responses were significantly larger for the location-script boundaries, suggesting that the “salience” of boundaries might vary across scripts ^70^. We also observed hippocampal activity increases at moments when key episodic details were being presented during events. This provides an interesting counterpart to prior work showing that subsequent recall and neural reinstatement are associated with hippocampal activity during events that is relatively low ^5^ and less correlated with the neocortex ^71^. Taken together, this suggests that the hippocampus is relatively disengaged within events, transiently engages to capture critical episodic details, and then encodes events into long-term memory at event boundaries ^5, 68^, possibly through sharp-wave ripple activity ^67, 72^.

### Changes in detail memory and event boundaries with script priming

In both our online study and our fMRI study, script priming influenced the story-specific details that participants could later recall, such that participants were significantly more accurate at responding to questions about details that were related to their attended script. This replicates classic work on schema priming, showing that an activated schema can help direct attention to relevant information and provide an organizational scaffold for forming memories ^40^. Although schemas can also play a role during recall by providing retrieval cues ^73^, in our design we prompted participants with the specific questions from both scripts during the memory test; the observed differences are therefore likely driven only by encoding-time processes.

We found that biasing attention to the location scripts in a multi-script narrative also caused significant shifts in subjective event boundaries. Participants who were location primed had segmentations that were significantly more aligned to the moments in the narratives where there was a transition in the location-script event sequence. Additionally, location primed participants segmented the story more similarly to each other compared to random pairs of participants. These results are consistent with previous work showing that an attentional focus on spatial information can impact annotated boundaries ^74^, demonstrating that this shift can also be driven by multi-stage location scripts.

Our results did not show a significant impact of social-script priming on annotated boundaries, and other work using the classic “burglar” and “home buyer” social scripts ^73^ similarly did not find evidence for greater alignment between annotated boundaries and prime-relevant story boundaries ^75^. This suggests that boundaries may be particularly sensitive to attentional manipulations of spatial or location-related information, though the reasons for this are unclear. One possibility is that the event boundaries in location scripts tend to be unambiguous (e.g. entering the checkout line at the grocery store) and therefore easier for participants to consistently align to, compared to event shifts in social situations which could be more gradual or uncertain. Another possibility is that there is a default orientation toward social events which occurs across all priming conditions, and the script priming therefore does not increase the salience of these socially-relevant boundaries. This second possibility is supported by our finding (in both the online and fMRI experiments) that participants performed better in general when answering socially-relevant questions, across priming conditions. Finally, this difference in priming impacts may not be about location vs social content per se, but other dimensions that vary between our two script categories, such as the degree to which they evoke strong emotions or the proportion of story content relevant to each script type.

### Shifting neural event boundaries through top-down script activation

Previous work using naturalistic stimuli has shown that attentional priming can impact representations during encoding. Priming participants with perspectives to elicit a social schema or a non-social schema impacted activation in angular gyrus and PHC ^41^, and manipulating participant’s initial beliefs before presenting an ambiguous story led to interpretation-specific responses in the DMN (including mPFC and angular gyrus) ^42^. Our findings extend this work by including a larger variety of priming conditions (eight possible scripts, as well as a no-prime baseline), and, critically, by using multi-stage temporal scripts rather than static perspectives. Using a Hidden Markov Model to identify neural transitions between stable states of activity ^5^, we found that location-script priming caused shifts in neural event boundaries that mirrored those seen in our online experiment. Location-primed participants exhibited boundaries that were significantly more aligned to location-relevant than socially-relevant narrative boundaries in mPFC (with similar, but more limited, effects in angular gyrus and PHC). These results provide the first evidence that activating an event script can have a top-down impact on the timing of neural pattern shifts in the DMN.

Event boundaries are often thought of as inherent features of a stimulus, and therefore predictable from the stimulus content alone ^76, 77^. For example, events are described as arising from “contextual stability in perceptual features” ^78^, “the spatiotemporal characteristics of the environment” ^79^, or “how fast various aspects of information from the environment tend to evolve” ^39^. Although the dynamics of the external environment are certainly a primary input to the event segmentation process, our findings instead favor a view of event boundaries as actively constructed in the mind and dependent on the prior knowledge and current goals of an individual ^13, 80^. Rather than simply inheriting the temporal dynamics of the environment, event segmentation is a cognitive process that optimizes the organization of continuous experience into the most relevant units. Individual differences in the segmentation process can have impacts on how much we remember ^81^, the order of our memories ^82^, and the contents of our memories ^83^.

Our findings shed new light on how event representations are constructed in the Default Mode Network from a broad set of prior knowledge, and how the activation of a schema for an event sequence can impact neural responses, perceived event boundaries, and subsequent memory for details. Our unique stimulus set allowed us to investigate how overlapping scripts can interact to support narrative perception and how attention can restructure the temporal dynamics of mental representations. These results identify mechanisms by which past experiences, distilled into schematic event scripts, change the way that we construct our present perceptions for realistic experiences.

## Acknowledgments

Thank you to Will Reblando for assistance in graphical design of the figures, Max Rosmarin for providing the voice recordings for the story stimuli, and to all the members of the Dynamic Perception and Memory Lab and Aly Lab for their suggestions and support. This work was supported by a Columbia University Lenfest Junior Faculty Development Award, to C.B.

## Author contributions

Alexandra De Soares: Conceptualization, Data curation, Investigation, Visualization, Software, Writing - original draft

Tony Kim: Data curation, Investigation, Writing - review & editing

Franck Mugisho: Data curation, Investigation, Writing - review & editing

Elen Zhu: Data curation, Investigation, Writing - review & editing

Allison Lin: Data curation, Investigation, Writing - review & editing

Chen Zheng: Data curation, Investigation, Writing - review & editing

Christopher Baldassano: Conceptualization, Funding acquisition, Methodology, Project Administration, Supervision, Visualization, Software, Writing - review & editing

## Supplemental Information

**Supplemental Figure S1:**
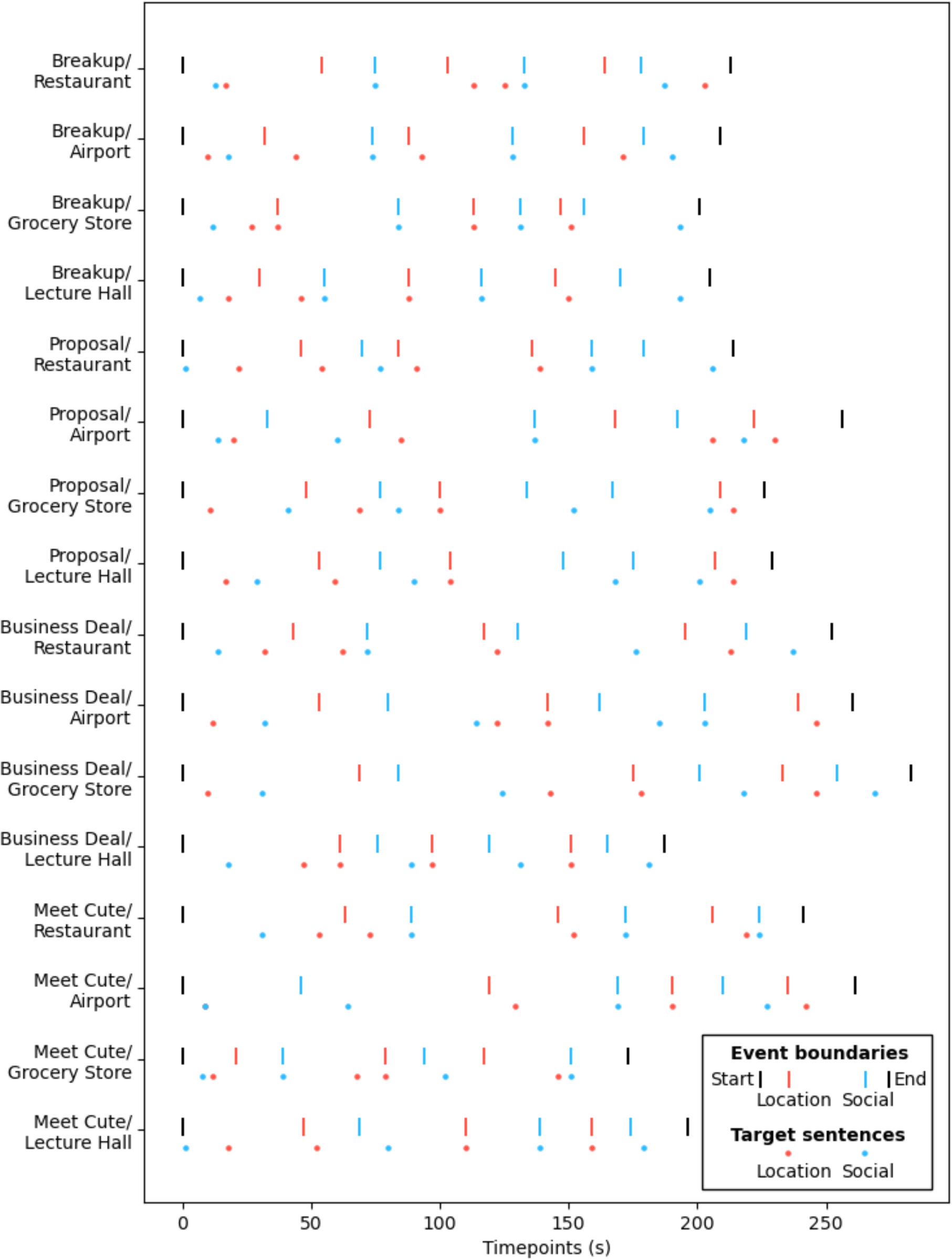
Timing of event boundaries and target sentences in each story, Related to Figure 1. Colored lines indicate the duration of each story and the timing of the three location-script boundaries and three social-script boundaries. Dots below the lines indicate the timing of target sentences containing episodic details probed during the memory test.

**Supplemental Figure S2:**
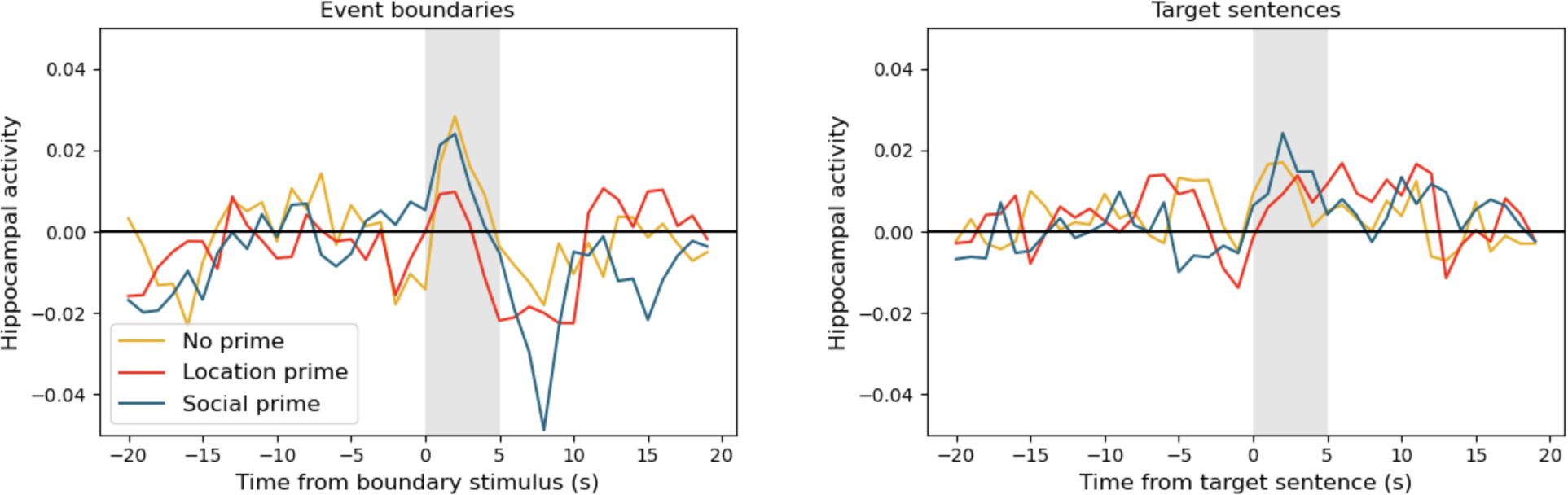
Hippocampal activity for boundary and target sentences, Related to Figure 2. We tested whether the univariate hippocampal activity (averaged across all hippocampal voxels) was related to event boundaries or to target sentences (which contained the answers to script-relevant questions). We z-scored the hippocampal timecourse for each story for each participant, and then extracted the timecourse values around event boundaries and target sentences. We defined the hippocampal response to a boundary/target to be the average activity in the 5 seconds following the boundary/target (gray band), and ran both a t-test for whether these responses were greater than zero (i.e. the mean response in the story) overall, and a repeated-measures ANOVA, with within-participant factors of priming condition and boundary/target script type. We found that there was a significant boundary-related response (t_35_=1.88, p=0.034) that was significantly larger for location boundaries (F_1,35_=7.37, p=0.010), but no main effect of priming (F_2,70_=1.49, p=0.232) or interaction between priming and boundary script type (F_2,70_=0.77, p=0.469). There was also a significant response to target sentences (t_35_=3.62, p<0.001), with no main effects of priming (F_2,70_=0.24, p=0.785) or script type (F_1,35_=0.05, p=0.830) or an interaction (F_2,70_=0.56, p=0.5728).

**Supplemental Figure S3:**
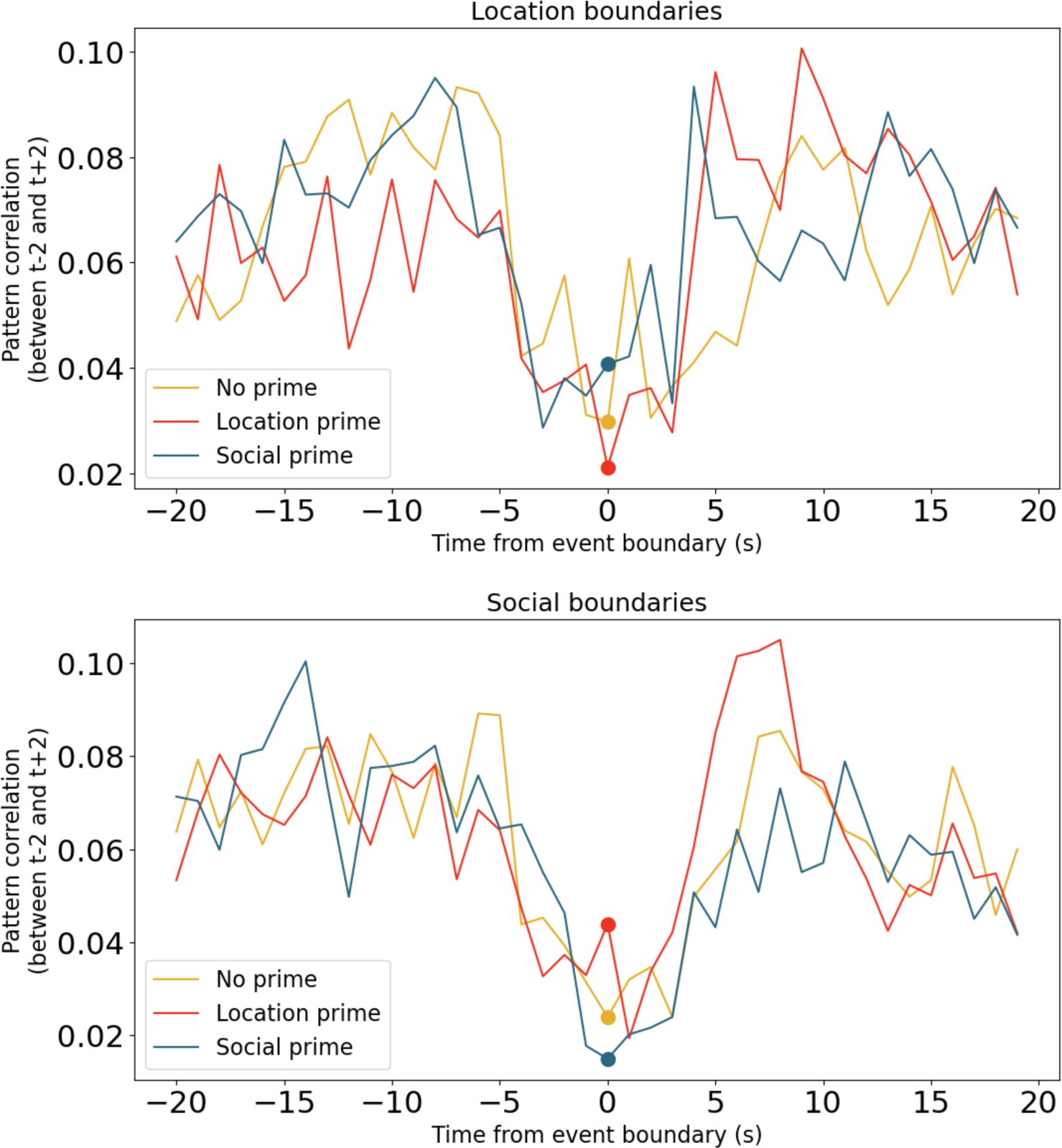
Pattern shifts in mPFC at event boundaries by priming condition, Related to Figure 5. Rather than measuring event change using the HMM, a simpler approach is to test whether voxel patterns are shifting more quickly around event boundaries than at other points in the story. For each timepoint *t* around a location (top) or social (bottom) boundary, we computed the correlation between the mPFC voxel pattern 2 seconds before *t* and 2 seconds after *t*, with the correlation across the boundary indicated by *t=0* (colored dots). The drop in correlation around event boundaries indicates shifts in mPFC patterns at boundary timepoints. Using the same statistical approach as in the main text (applied to pattern correlations rather than HMM event change), we find only marginal evidence for an overall priming effect (p=0.097), but do find significant alignment (i.e. reduced pattern correlation) to location boundaries with location priming (p=0.022), and significant alignment to social boundaries with no priming (p=0.037) or social priming (p=0.006), which are significantly different from location priming (p=0.031).

**Supplemental Figure S4:**
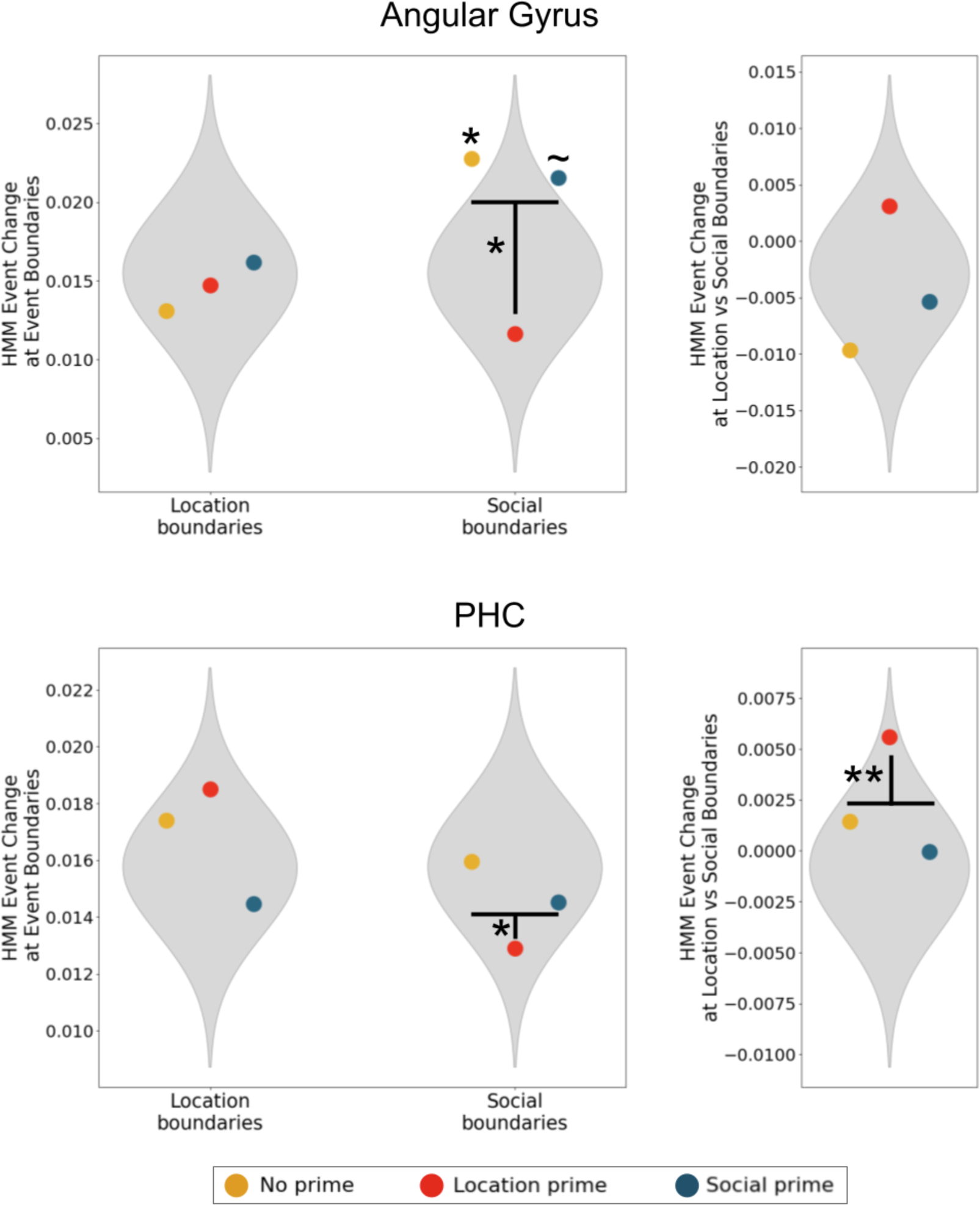
HMM-boundary alignment for angular gyrus and PHC, Related to Figure 5. Applying the same analysis as in main text Figure 5c to angular gyrus and PHC, we observe a significant overall effect of location priming in shifting alignment from social to location boundaries in PHC (p=0.008; angular gyrus p=0.126). We did not observe significant alignment to location boundaries in any priming condition in either region (all p>0.10) but did find significant decreases in alignment to social boundaries in the location-primed group for both regions (angular gyrus p=0.030; PHC p=0.018). In angular gyrus the social boundaries were aligned to neural boundaries significantly for the no-prime group (p=0.042; PHC p=0.468) and marginally for the social-prime group (p=0.075; PHC p=0.703). ∼ p < 0.10, * p < 0.05, ** p < 0.01

**Supplemental Figure S5:**
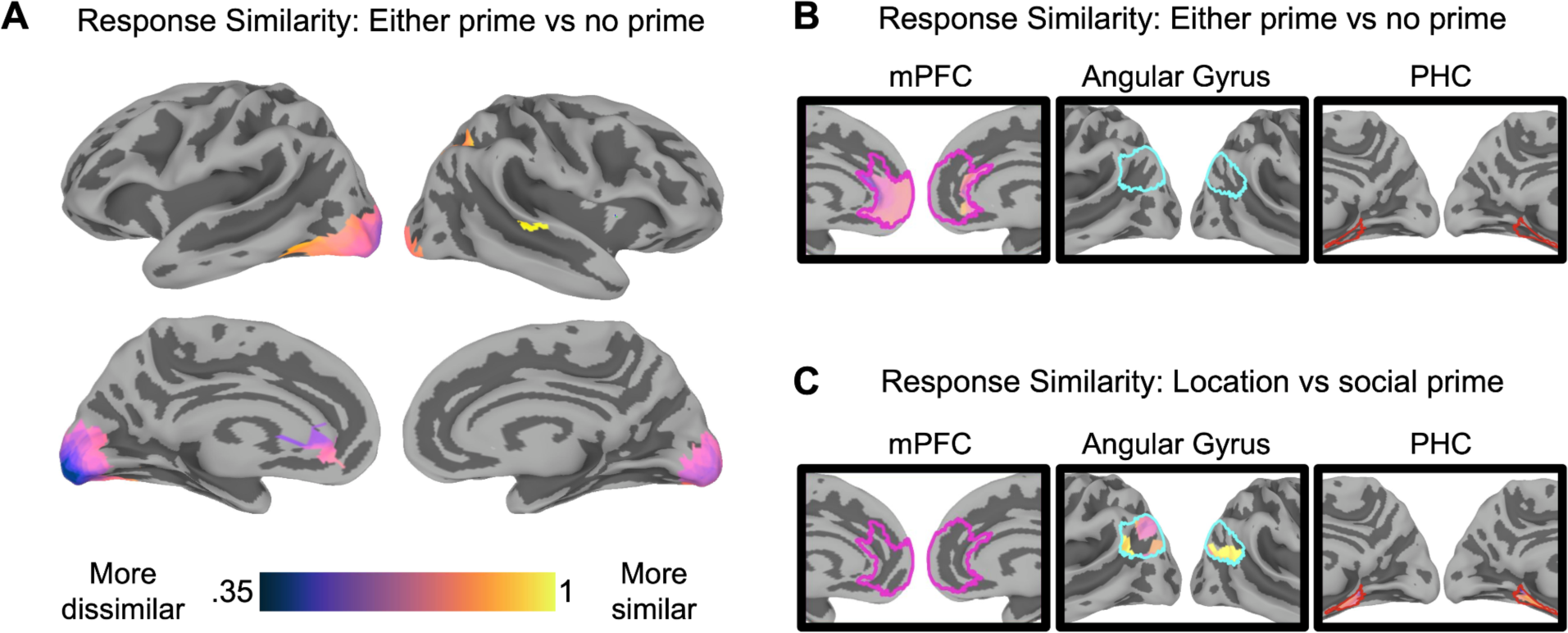
Neural activation during story listening differs with prime, Related to Figure 5. We conducted a model-free exploratory analysis to test for any systematic changes in neural responses across priming groups. Unlike our main HMM analysis, which provided a targeted test specifically for shifts in event boundaries, this analysis allows us to identify brain regions whose responses were strongly impacted by attentional priming in any way, regardless of whether the change is related to event boundaries. Note that this more exploratory approach has more limited statistical power and does not test a specific cognitive hypothesis about how priming impacts neural dynamics. For each story, we randomly sampled two groups of 4 participants from each priming condition, and correlated the average response timecourses between all pairs of these six groups. If priming produces a systematic shift in responses, two groups with the same prime should have higher correlations than groups with different primes. We mathematically compared these values using a Response Similarity approach derived in our previous work ^110^; specifically, we divide the across-group correlation by the geometric mean of the within-group correlations. If this ratio is significantly smaller than 1, this indicates that there are systematic differences between the groups’ mean responses. We performed permutation tests by shuffling the priming conditions of the participants in a story 1,000 times (for computational efficiency, we first performed only 100 permutations in each searchlight, and then performed the rest of the permutations only if at least 95% of the null Response Similarities were less than the real Response Similarity). For each vertex, we averaged these values across all searchlights that included this vertex, and computed a p value as the fraction of the 1,001 Response Similarity values (including the real value) that were at least as small as the real value. These were converted to *q* values using the false discovery rate in AFNI ^108^, across the full cortical surface or as a small volume correction within *a priori* ROIs (mPFC, PHC, angular gyrus). (A) Comparing the no-prime group to the average of both priming groups, we identified numerous areas that showed significant differences (significantly diminished Response Similarity) including regions in the intraparietal sulcus, the MTC, mPFC, and early visual cortex. (B) We additionally performed a small volume correction to look for an effect specifically in our three regions of interest. We found that there were significant differences in Response Similarity for primed vs. unprimed participants only in the mPFC. (C) Computing the Response Similarity between the participants who received location vs. social priming, no voxels in the searchlight analysis survived FDR correction when considering the whole cortical surface, but after performing a small volume correction in our three regions of interest we observed significant effects within angular gyrus and PHC. All maps are thresholded at q<0.05.

**Supplemental Figure S6:**
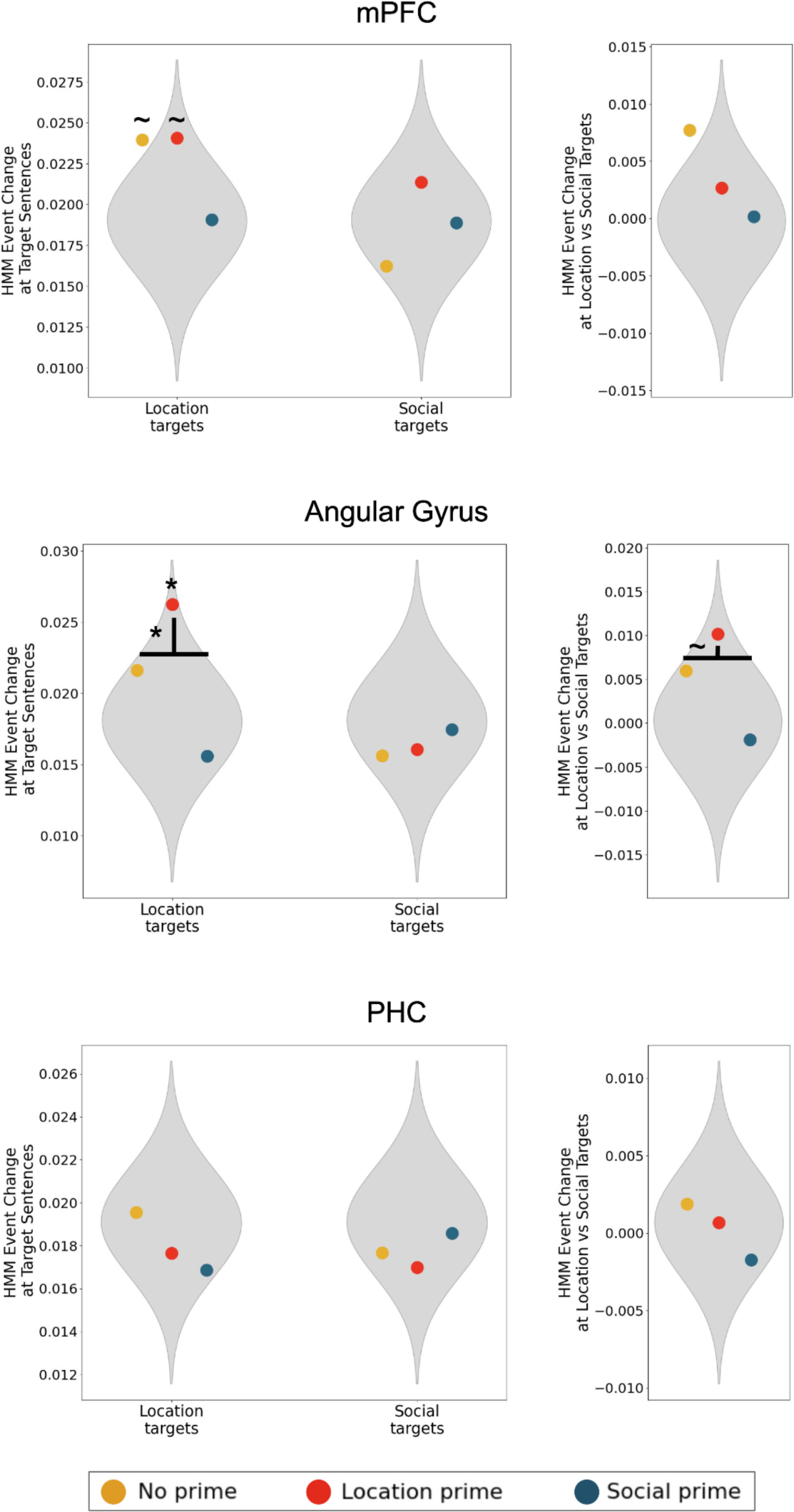
HMM event change at target sentences, Related to Figure 5. We repeated our HMM fMRI analysis, but measured the degree of HMM boundary shift at target sentences rather than at our putative event boundaries. We did not observe significant overall effects of priming in any of our ROIs (mPFC p=0.94; angular gyrus p=0.098; PHC p=0.35). Running exploratory tests for alignment to location and social boundaries within each ROI, there were no significant alignment or priming effects in mPFC or PHC; in angular gyrus the only significant alignment was for location-primed participants to location-related targets (p=0.02), which was significantly higher than for other priming conditions (p=0.03). ∼ p < 0.10, * p < 0.05

**Supplemental Table S1:** Timing statistics for the event durations for both script types, Related to STAR Methods.

|  | Event 1 | Event 2 | Event 3 | Event 4 |
| --- | --- | --- | --- | --- |
| ***Location event durations*** | 53.06 ± 21.85 | 65.94 ± 19.49 | 63.56 ± 21.59 | 42.81 ± 16.4 |
| ***Social event durations*** | 68.75 ± 16.08 | 73.25 ± 23.66 | 45.25 ± 15.88 | 38.12 ± 14.49 |

**Supplemental Table S2:** Timing statistics for the target sentences for both script types, Related to STAR Methods.

|  |  | Question 1 | Question 2 | Question 3 | Question 4 |
| --- | --- | --- | --- | --- | --- |
| ***Location target sentences*** | **Time in story (s)** | 20.94 ± 12.64 | 76.06 ± 31.82 | 124.38 ± 37.5 | 193.38 ± 38.39 |
|  | **Time within event (s)** | 20.94 ± 12.64 | 23.0 ± 23.89 | 5.38 ± 10.03 | 10.81 ± 10.2 |
| ***Social target sentences*** | **Time in story (s)** | 17.44 +- 11.6 | 79.38 +- 20.12 | 151.0 +- 28.51 | 204.0 +- 26.39 |
|  | **Time within event (s)** | 17.44 +- 11.6 | 10.62 +- 12.65 | 9.0 +- 12.67 | 16.75 +- 11.87 |

**Supplemental Table S3:** Jaccard similarity of behavioral event responses to event boundaries and target sentences, Related to Figure 4. We reran our behavioral analyses of the online participant data (in which participants were asked to mark when the next “part” of the story began), but now measured Jaccard similarity to the target sentences rather than event boundaries. Overall we found significantly less alignment to target sentences than to our putative event boundaries, for location-related targets/boundaries in all three priming conditions (no-prime: t_100_=3.28, p=0.001; location-prime: t_105_=3.70, p<0.001; social-prime: t_100_=3.04, p=0.003) and for social-related targets/boundaries in all three priming conditions (no-prime: t_100_=3.83, p<0.001; location-prime: t_105_=5.04, p<0.001; social-prime: t_100_=2.53, p=0.013). Interestingly, although the frequency of responses at target sentences was relatively low compared to event boundaries, we did observe priming effects at target sentences; alignment was significantly different across priming conditions for location targets (F_2,305_ = 4.24, p = 0.015; location priming significantly higher than social priming, p = 0.013 by post-hoc Tukey test) and social targets (F_2,305_ = 3.21, p = 0.042; no significant differences in post-hoc Tukey test).

|  | No Prime | Location Prime | Social Prime |
| --- | --- | --- | --- |
| **Location Boundaries** | 0.197 | 0.226 | 0.173 |
| **Location Targets** | 0.141 | 0.171 | 0.125 |
| **Social Boundaries** | 0.162 | 0.174 | 0.175 |
| **Social Targets** | 0.114 | 0.109 | 0.143 |

Table S4. Stimulus sentences and timing information.

## STAR★Methods

### Resource availability

#### Lead contact

Further information and requests for resources should be directed to and will be fulfilled by the lead contact, Christopher Baldassano.

#### Materials availability

The stories developed for this study have been included as Supplemental Table S4, and the audio versions of the stories are available at https://figshare.com/projects/Script_Combination_Stories/168656.

#### Data and code availability

Data from the fMRI experiment is available in BIDS format at https://doi.org/10.18112/openneuro.ds004631.v1.0.0. Analysis code and anonymized data from the behavioral experiment are available at https://github.com/dpmlab/ScriptPriming [a DOI will be generated for the final version of the repository upon paper acceptance].

### Experimental model and study participant details

#### fMRI experiment

We collected data from a total of 38 subjects (21 female; ages 19 - 32 years; 22 white, 11 asian, 2 black, 2 mixed; 7 Hispanic or Latino). All participants were fluent in English, and 32 were native English speakers. All participants had normal or corrected-to-normal vision. The experimental protocol was approved by the Institutional Review Board of Columbia University, and all participants gave their written informed consent. Two participants were excluded due to excessive motion in the scanner (more than 60% of recall timepoints identified as motion outliers, as described below), yielding a final sample of 36 (our target sample size). This sample size was chosen based on prior experiments examining event boundaries with fMRI ^25, 84^ and to ensure that each of the 36 possible combinations of priming conditions (see “Experimental design” below) was assigned to one participant.

#### Online behavioral experiment

Data were collected online using Prolific (www.prolific.co). We collected data from 387 participants (181 females, age 18 - 75) who were 96% native English speaking and we compensated them $3.17- $5. We excluded 10 participants who did not answer the short answer questions following the story, resulting in a final sample of n=377. For the short-answer analyses, we additionally removed 2 participants whose short-answer data was not correctly recorded (yielding n=375), and for the event-boundary analyses, we additionally excluded 28 participants who did not mark any event boundaries and 41 participants who said that more than half of the sentences were event boundaries (yielding n=308). This sample size was chosen to ensure that at least 5 participants were assigned to each of the 3 possible primings of each of the 16 stories. The experimental protocol was approved by the Institutional Review Board of Columbia University.

### Method details

#### fMRI experiment

##### Stimuli

Each narrative stimulus included events reflecting one “location script” and one “social script”. Location scripts were all common sequences of activities that occur in a specific kind of place to accomplish a specific goal: eating at a restaurant, catching a flight at an airport, shopping at a grocery store, and attending a lecture. Social schemas were all common sequences of events for changes in relationships and were not constrained to a particular location: initiating a breakup, proposing marriage, conducting a business deal, and experiencing a “meet cute”. We expected participants to be familiar with these schemas either from their everyday experiences or from media exposure. Each script consisted of a temporal sequence of four events (see Table 1).

**Table 1:**
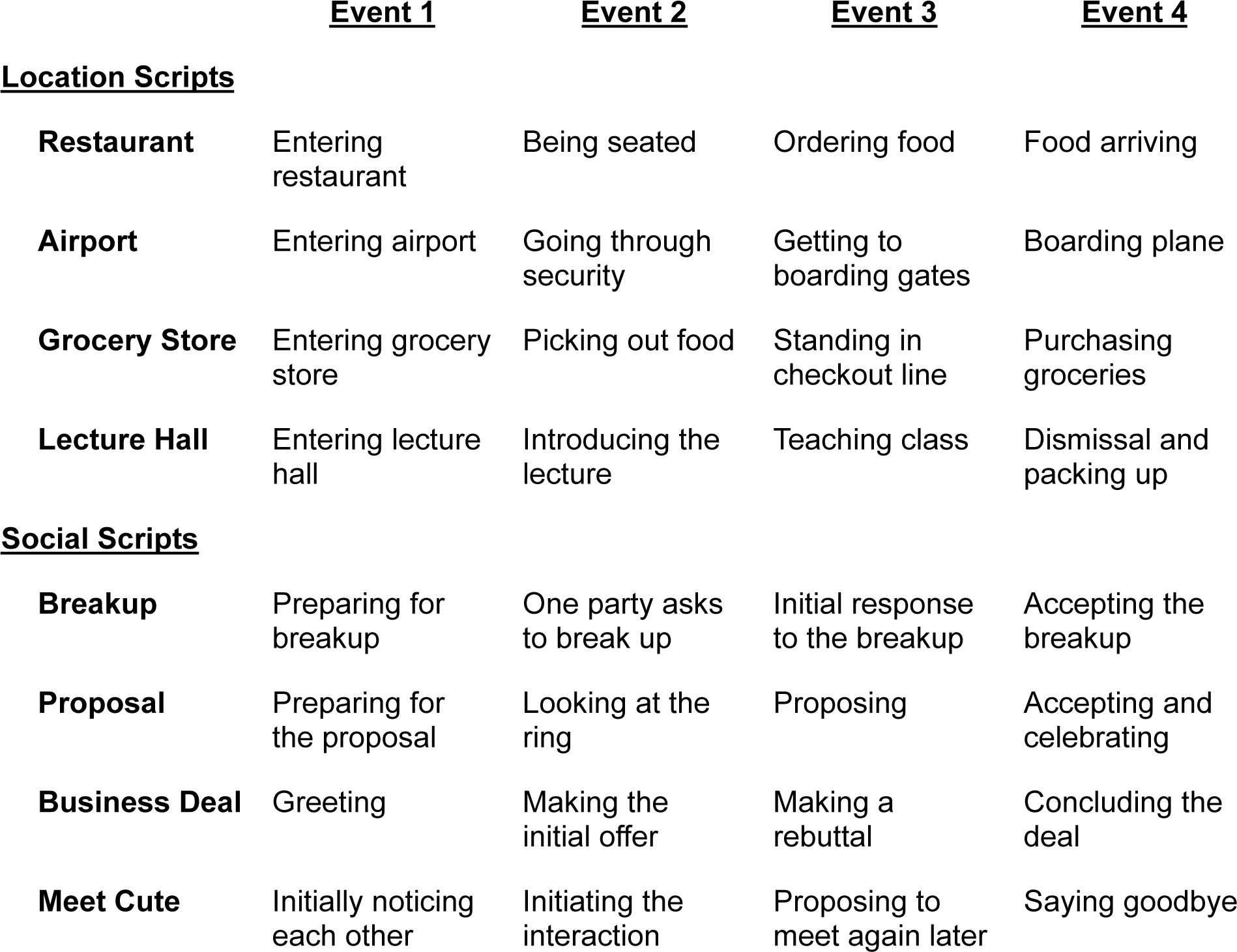
Location and social scripts and their events. Each location and social script has a stereotyped four-event sequence which was consistent in each story that contained that script.

We created 16 custom stories for this study, each consisting of a unique combination of one of the four location scripts and one of the four social scripts. Each story proceeded through the four events of each script in parallel. Each event lasted 56.3 seconds on average, but the duration (and number of sentences) for each event varied across stories. Each story consisted of 27-41 sentences (average of 33.19 ± 3.88), and each sentence had a duration of 6.82 ± 2.74 seconds. The mean and standard deviations of the four events for each script type are provided in Supplemental Table S1. An ANOVA with factors of event number (1-4) and specific script (e.g. Restaurant) found significant variation across event number (location events: F_3,48_=4.07, p=0.011; social events: F3,48=14.86, p<0.001) but no effect of script on event length (location events: F_3,48_=0.70, p=0.553; social events: F_3,48_=0.87, p=0.464) or an interaction between event number and script (location events: F_9,48_=0.83, p=0.591; social events: F_9,48_=1.47, p=0.186), indicating that there were no significant differences between scripts in the temporal structure of their events.

To allow us to behaviorally and neurally distinguish between responses related to location event boundaries versus social event boundaries, the event onsets for the two scripts were interleaved and never coincided in the same sentence (except for the very first sentence, which started the first event for both scripts). Each story had a unique set of characters and a unique setting, such that the only similarities between stories came from overlap in their event scripts. Stories were presented as audio narratives (read by the same professional voice actor) and were on average 3 minutes and 45 seconds long. While participants listened to the story they were presented with a still screen displaying the title of the story and the pictures (from the 10k US Adult Faces Database ^85^) and names of the two main characters for that story.

##### Script priming

In order to prime participants, we asked them to take on a perspective pertaining to the script, inspired by previous work on schema priming ^40, 41, 73^. To encourage attention to the specific four-event sequence of each script, we provided participants with a sequence of four script-specific questions they would need to answer at the end of the story. Each of these questions corresponded to one of the script events, as shown in Table 2. Although we expected that all participants would be familiar with the eight scripts in our stories, this priming procedure was intended to provide a standardized definition of the schematic events for each script and what should be considered a script-relevant detail. In each story there were eight sentences, which we refer to as “target sentences,” which contained details relevant to either the location or social questions. Descriptive statistics about the timing of target sentences are given in Supplemental Table S2.

**Table 2:** Location and social scripts questions. Priming for each location and social script involved learning a sequence of four questions that corresponded to each event in the script.

|  | <u>Question 1</u> | <u>Question 2</u> | <u>Question 3</u> | <u>Question 4</u> |
| --- | --- | --- | --- | --- |
| **<u>Location Scripts</u>** |  |  |  |  |
| **Restaurant** | How is the restaurant decorated? | What are the menus like? | What does each person order? | How do the clients like the food? |
| **Airport** | When the clients arrive at the airport, how much time do they have until their flight departs? | What is the reason for the hold-up at security? | Toward which gate are the clients walking? | What section and seat does each client sit in on the plane? |
| **Grocery Store** | What is the grocery store like upon entering? | What items do the clients pick out to buy? | How many checkout lanes are open and which one do the clients step into? | How much are the groceries and what method of payment do the clients use? |
| **Lecture Hall** | What is the lecture hall like? | What class are the students in and what is the day's lecture about? | What is something taught in lecture? | What is the next assessment/ assignment for the class, and when is it scheduled/due? |
| **<u>Social Scripts</u>** |  |  |  |  |
| **Breakup** | For how long has the initiator of the breakup been thinking about breaking up with his/her partner? | What is the initial reason stated by the initiator for why he/she is breaking up? | Does the person who is being broken up with want to break up, and what's the reason stated by them for this? | Who wants what items back as a result of the breakup? |
| **Proposal** | For how long has the couple been dating? | How many diamonds are on the ring, and what is the diamond color? | In/on what item is the ring presented? | Who does the new fiancée text first? |
| **Business Deal** | What is each business person's job title? | How much money is at stake in the initial offer made? | What is the name of the other industry competitor? | With what gesture do the business partners secure the deal? |
| **Meet Cute** | What object is the initiator of the interaction holding when he/she first notices the other person? | What is the initial question that begins the conversation between the couple? | When will be the next time the couple meets and for what occasion? | What time is it when the couple parts? |

Participants were primed with two of the location scripts and two of the social scripts. Before the scanning session, each participant was shown the four perspectives they would be asked to take on. Then for each perspective, they were taught the four script-specific questions. Participants then had to correctly identify and order the questions for each script from six possible answers, with the correct sequence being re-shown after each attempt. After completing the training on all four perspectives, they were asked to reorder the questions for each perspective again without being shown the correct sequence. When participants were in the scanner and about to listen to a story relevant to one of their primes, they would be told which perspective to take on, and had to correctly identify and order the four script-specific questions before moving onto the story.

##### Data collection and processing

Data were collected on a 3T Siemens Prisma scanner with a 64-channel head/neck coil. Functional images were obtained with an interleaved multiband EPI sequence (TE = 30 ms, flip angle = 62°, multiband = 3, FOV = 205 mm x 205 mm, 48 oblique axial slices), resulting in a voxel size of 2.5 mm isotropic and a TR of 1s. Whole-brain high resolution (1.0 mm isotropic) T-1 weighted structural images were acquired with an MPRAGE sequence.

Results included in this manuscript come from preprocessing performed using *fMRIPrep* 1.5.6 ^86, 87^ (RRID:SCR_016216), which is based on *Nipype* 1.4.0 ^88, 89^ (RRID:SCR_002502).

###### Anatomical data preprocessing

The T1-weighted (T1w) image was corrected for intensity non-uniformity (INU) with N4BiasFieldCorrection ^90^, distributed with ANTs 2.2.0 ^91^ (RRID:SCR_004757), and used as T1w-reference throughout the workflow. The T1w-reference was then skull-stripped with a *Nipype* implementation of the antsBrainExtraction.sh workflow (from ANTs), using OASIS30ANTs as target template. Brain tissue segmentation of cerebrospinal fluid (CSF), white-matter (WM) and gray-matter (GM) was performed on the brain-extracted T1w using fast (FSL 5.0.9, RRID:SCR_002823) ^92^. Brain surfaces were reconstructed using recon-all (FreeSurfer 6.0.1, RRID:SCR_001847) ^93^, and the brain mask estimated previously was refined with a custom variation of the method to reconcile ANTs-derived and FreeSurfer-derived segmentations of the cortical gray-matter of Mindboggle (RRID:SCR_002438) ^94^. Volume-based spatial normalization to one standard space (MNI152NLin2009cAsym) was performed through nonlinear registration with antsRegistration (ANTs 2.2.0), using brain-extracted versions of both T1w reference and the T1w template. The following template was selected for spatial normalization: *ICBM 152 Nonlinear Asymmetrical template version 2009c* ^95^ (RRID:SCR_008796; TemplateFlow ID: MNI152NLin2009cAsym).

###### Functional data preprocessing

For each BOLD run (across all tasks and sessions), the following preprocessing was performed. First, a reference volume and its skull-stripped version were generated using a custom methodology of *fMRIPrep*. A deformation field to correct for susceptibility distortions was estimated based on *fMRIPrep*’s *fieldmap-less* approach. The deformation field is that resulting from co-registering the BOLD reference to the same-subject T1w-reference with its intensity inverted ^96, 97^. Registration is performed with antsRegistration (ANTs 2.2.0), and the process regularized by constraining deformation to be nonzero only along the phase-encoding direction, and modulated with an average fieldmap template ^98^. Based on the estimated susceptibility distortion, a corrected EPI (echo-planar imaging) reference was calculated for a more accurate co-registration with the anatomical reference. The BOLD reference was then co-registered to the T1w reference using bbregister (FreeSurfer) which implements boundary-based registration ^99^. Co-registration was configured with six degrees of freedom. Head-motion parameters with respect to the BOLD reference (transformation matrices, and six corresponding rotation and translation parameters) are estimated before any spatiotemporal filtering using mcflirt (FSL 5.0.9) ^100^. The BOLD time-series were resampled to surfaces on the following spaces: *fsaverage6*. The BOLD time-series (including slice-timing correction when applied) were resampled onto their original, native space by applying a single, composite transform to correct for head-motion and susceptibility distortions. These resampled BOLD time-series will be referred to as *preprocessed BOLD in original space*, or just *preprocessed BOLD*. The BOLD time-series were resampled into standard space, generating a *preprocessed BOLD run in [‘MNI152NLin2009cAsym’] space*. First, a reference volume and its skull-stripped version were generated using a custom methodology of *fMRIPrep*. Several confounding time-series were calculated based on the *preprocessed BOLD*: framewise displacement (FD), DVARS and three region-wise global signals. FD and DVARS are calculated for each functional run, both using their implementations in *Nipype* ^101^. The three global signals are extracted within the CSF, the WM, and the whole-brain masks. The head-motion estimates calculated in the correction step were also placed within the corresponding confounds file. The confound time series derived from head motion estimates and global signals were expanded with the inclusion of temporal derivatives for each ^102^. Frames that exceeded a threshold of 0.5 mm FD or 1.5 standardised DVARS were annotated as motion outliers. All resamplings can be performed with *a single interpolation step* by composing all the pertinent transformations (i.e. head-motion transform matrices, susceptibility distortion correction when available, and co-registrations to anatomical and output spaces). Gridded (volumetric) resamplings were performed using antsApplyTransforms (ANTs), configured with Lanczos interpolation to minimize the smoothing effects of other kernels ^103^. Non-gridded (surface) resamplings were performed using mri_vol2surf (FreeSurfer).

Many internal operations of *fMRIPrep* use *Nilearn* 0.6.1 ^104^ (RRID:SCR_001362), mostly within the functional processing workflow. For more details of the pipeline, see the section corresponding to workflows in *fMRIPrep*’s documentation.

After *fMRIPrep*, the data (now in fsaverage6 and MNI152 space) was further preprocessed by a custom python script that: removed from the data (via linear regression) any variance related to the six degrees of freedom motion correction estimate and their derivatives, mean signals in the CSF and white matter, motion outlier timepoints (defined above), and a cosine basis set for high-pass filtering w/ 0.008 Hz (125s) cut-off; and z scored each run to have a zero mean and SD of 1. We divided the runs into the portions corresponding to each stimulus, extracting timepoints from the start to the end of the stimulus presentation shifted by 5 seconds to account for hemodynamic lag. All subsequent analyses, described below, were performed using custom python scripts and the Brain Imaging Analysis Kit (http://brainiak.org/) ^105^.

##### ROI and searchlight definition

We used ROIs in the default mode network previously found to be responsive to schematic content ^25^: angular gyrus (1868 vertices), middle temporal cortex (MTC, 2118 vertices; labeled in the previous study as superior temporal sulcus, STS), superior frontal gyrus (SFG; 2461 vertices), medial prefrontal cortex (mPFC; 2069 vertices), parahippocampal cortex (882 vertices), and posterior medial cortex (PMC; 2495 vertices). These ROIs were originally derived from a resting-state network atlas on the fsaverage6 surface ^106^. We also used, as a control region, a previously-defined auditory cortex region (1589 vertices), which we expected to be largely insensitive to semantic properties of narratives ^25^. Additionally, to extract the hippocampus as an ROI, we used the freesurfer subcortical parcellation provided in the *fmriprep* outputs, and included voxels which were common to at least 80% of participants (785 voxels).

Searchlight ROIs were defined as circular regions on the cortical surface, by identifying all vertices within 11 edges of a center vertex along the fsaverage6 mesh. Since the average edge length between vertices is 1.4mm, searchlights had a radius of approximately 15mm. We defined a circular searchlight around every vertex on a hemisphere, and then iteratively removed the most redundant searchlights (i.e. those whose vertices were covered by the most other searchlights). We stopped removing searchlights when doing so would cause some vertices to be covered by fewer than six searchlights. This yielded approximately 1000 searchlights on each hemisphere.

##### Experimental design

We first tested the audio presentation and recording equipment by playing a short audio clip (unrelated to the narratives) to verify that the volume level was set correctly and by asking participants to talk about their breakfast that day into the microphone. Subjects were then presented with 6 stories in each of 2 experimental runs, using PsychoPy (RRID:SCR_006571)^107^. Participants were each trained on two of the four location scripts and two of the four social scripts, using the procedure described above; this produced C(4,2)*C(4,2)=36 possible priming conditions, with each condition assigned to a different participant. Each participant listened in total to 12 stories: 4 stories that used the primed location scripts but not the primed social scripts; 4 stories that used the primed social scripts but not the primed location scripts; and 4 stories that used neither of the primed location or social schemas. Participants did not hear the remaining 4 stories for which both the location and social scripts had been primed. Story order and priming were shuffled in each block. If a story contained one of the primed event scripts, participants were reminded before the story of the perspective they were supposed to take on and took a quiz to identify and correctly order the four script-relevant questions. Participants would not proceed to the story until they correctly ordered the questions. At the onset of the story, the story image would appear on a gray background followed by 5 seconds of silence. When the story finished, the story image would disappear followed by an additional 5 seconds of silence. Subjects were not explicitly told that they would be asked to recall the stories. After participants listened to the 12 stories, they were then asked to recall stories in the same order as they listened (data not reported in this paper).

After recall, participants were asked outside of the scanner to answer 8 short answer questions about each story. These questions were the script-specific questions for the social and location perspectives relevant to that story.

#### Online behavioral experiment

##### Stimuli

Stimuli were presented using Pavlovia (pavlovia.org). The stimuli in the online experiment were the same stories as in the fMRI experiment. However, rather than listening to the story continuously, participants listened to each sentence individually. When each sentence had finished, participants indicated if the just-presented sentence was the beginning of a new “part” of the story or not, the story then proceeded to the next sentence. After participants listened to the story and made judgments on all sentences, they performed a brief distractor task where they were asked to remember difficult-to-spell words, and then they were asked the same 8 script-specific questions for the two scripts relevant to the story in the same manner as in the fMRI study. Finally participants typed what they recalled about the story (data not analyzed in this paper), and filled out a demographics questionnaire about in Qualtrics (www.qualtrics.com).

##### Priming

Participants were randomly assigned to hear one story, and were randomly assigned to either be not primed, primed with the location script, or primed with the social script. Priming was done in the same manner as in the fMRI study, with participants asked to “take on” a perspective while listening to the story. Participants were first shown the four script-specific questions (Table 2) in the correct order, and were then given the quiz where they had to select the questions in the right order. If they made an incorrect choice they would be shown the correct sequence again and then had to re-take the quiz. Once they had correctly ordered all the questions, they would be asked to perform the ordering correctly a second time; if they made a mistake, they were re-presented with the quiz until they produced the correct ordering.

#### Quantification and statistical analysis

##### Short Answer Priming Effect Analysis

We scored the short answers of all participants (from both the fMRI and online studies) on a scale from 0-3; a score of 0 indicated a failure to respond or recall any accurate details, a score of 3 indicated perfect recall of all episodic details for a particular question, and scores of 1-2 denoted partial recall (according to a scoring rubric for each question). We then computed the average location question score by averaging the ratings of all location questions, and in the same way we computed an average social question score. We ran a mixed effects model separately for each experiment with prime, question type, and their interaction as fixed effects, and a subject-specific random intercept. We also tested for a significant difference in average score between priming groups by running a Welch independent-samples t-test between location-primed and social-primed groups for each question type.

##### Comparison to putative event boundaries

We computed the Jaccard similarity between each online participant’s segmentation and the putative location and social boundaries of each story. These putative boundaries were the sentences that were designed to indicate a shift to the next event in the location or social script (see Table 1). In order to test whether participants in each priming group had segmentations that aligned with these event boundaries, we computed the Jaccard similarity index between each participant’s segmentations and the putative boundaries from both scripts. We computed the average similarity in each priming group, and compared to a null in which between-boundary durations were preserved but the order of event durations was shuffled 10,000 times. We computed a p value as the fraction of agreements in the 10,001 null and real values that were at least as large as in the real agreement.

To test for differences in alignment between priming conditions for each type of event boundary, we used a one-way ANOVA to test for differences in mean agreement to the story boundaries between priming groups, and if there was a significant difference we ran Tukey’s test to see which means were significantly different.

##### Within-group similarity of event boundaries

In order to determine if priming caused online participants to have more similar event boundary judgments, we computed the Jaccard similarity index between all pairs of participants within each priming group for each story, and averaged across stories. Permutation testing was used for assessing statistical significance; the priming conditions for the participants in each story were shuffled 10,000 times and within-group similarity was computed for each null dataset, and the p value was computed as the fraction of the 10,001 null and real similarities that were at least as large as the real within-group similarity.

##### Location and Social Script Effect Analysis

To measure the extent to which event patterns generalized across stories with the same location script in a brain region, we first computed for each fMRI participant the average spatial activity pattern for each of the four events of each story (defined according to the putative location-related event boundaries). For all pairs of stories, we correlated all four event patterns in one story for one participant with the four event patterns in the second story averaged across all other participants. We averaged the four correlations between corresponding events (e.g. event 1 in story 1 with event 1 in story 2) and subtracted the average of all other event correlations as a baseline. We iteratively held out each participant, recomputing the 16 story x 16 story event correlation matrix for each held-out participant, and then averaged these matrices to produce a single event correlation matrix. To compute the location script effect, we compared the average event correlation for pairs of stories with shared location scripts to the event correlation for pairs of stories with no shared scripts. If the same-script correlations are larger, this indicates that the location script of a story influences the event patterns in a brain region. We then repeated this entire process for the social scripts, computing event patterns according to the social-relevant boundaries and comparing event similarity for stories with shared social scripts versus no shared scripts.

We used permutation testing to assess statistical significance. After computing the location and script effects, we repeated the process 1000 times, randomly shuffling the identities of the 16 stories each time. For our *a priori* ROIs, we computed p values directly as the fraction of the 1001 effect values (including the real value) that were at least as large as the real value. For the searchlight analysis, in order to better estimate small p values, we approximated the distribution of null values using a Normal distribution, and converted the real effect value into a z score relative to this distribution (i.e. the number of standard deviations by which the real value exceeded the mean of the null distribution). A z value was computed for each surface vertex as the average of z values from all searchlights that included that vertex. The vertex z values were then converted to p values using the survival function of the Normal distribution, and finally converted into a map of q values using the same false discovery rate correction that is used in AFNI ^108^.

##### Hidden Markov Model Analysis

In order to test if priming had an effect on neural event boundaries, we used a Hidden Markov Model (HMM) approach ^5^ to compare fMRI participants’ neural event boundaries to the putative location and social event boundaries in the narratives. For a given brain region and story, we first averaged the spatiotemporal pattern of brain activation within each of the three priming groups. We then fit an HMM with 4 events to the average pattern of activation of each group, producing a probabilistic estimate of which of the four events was active at each timepoint. We used this probability distribution to compute a measure of event change at each timepoint by taking the derivative of the expected value of the active event at each timepoint ^109, 110^. Note that event change effectively provides a continuous measure of “boundary-ness,” rather than defining discrete timepoints as HMM boundaries (e.g. by identifying timepoints at which the maximum-probability event changes). This provides a more sensitive approach for detecting subtle shifts in boundary dynamics, and does not require setting an arbitrary temporal distance threshold for computing matches between neural and behavioral boundaries.

To assess the match between the neural event shifts and the putative event boundaries of the location and social scripts in each story, we computed the average event change at the location boundaries and at the social boundaries for each story for each priming group. We constructed null boundary sets in which between-boundary durations were preserved but the order of event durations was shuffled; because there were 4 events, this resulted in 4!=24 boundary sets for each script for each story (1 original and 23 null). After averaging both the real alignment and null alignments across stories, we computed a p value for the alignment of each priming group to each boundary type by modeling the null distribution as a Normal distribution and then computing the area of this Normal that exceeded the real alignment value. Similarly, we tested whether there were significant differences in boundary alignment between the location and non-location priming groups by measuring the difference between the location priming group and the average of the social and no prime groups, comparing the difference for the real values to the differences in the null distribution. Finally, we computed an overall measure of our priming effect by computing the alignment to location boundaries minus social boundaries for each priming group, and testing whether the preference for location boundaries was larger for the location-primed group using the same statistical procedure.

### Key resources table

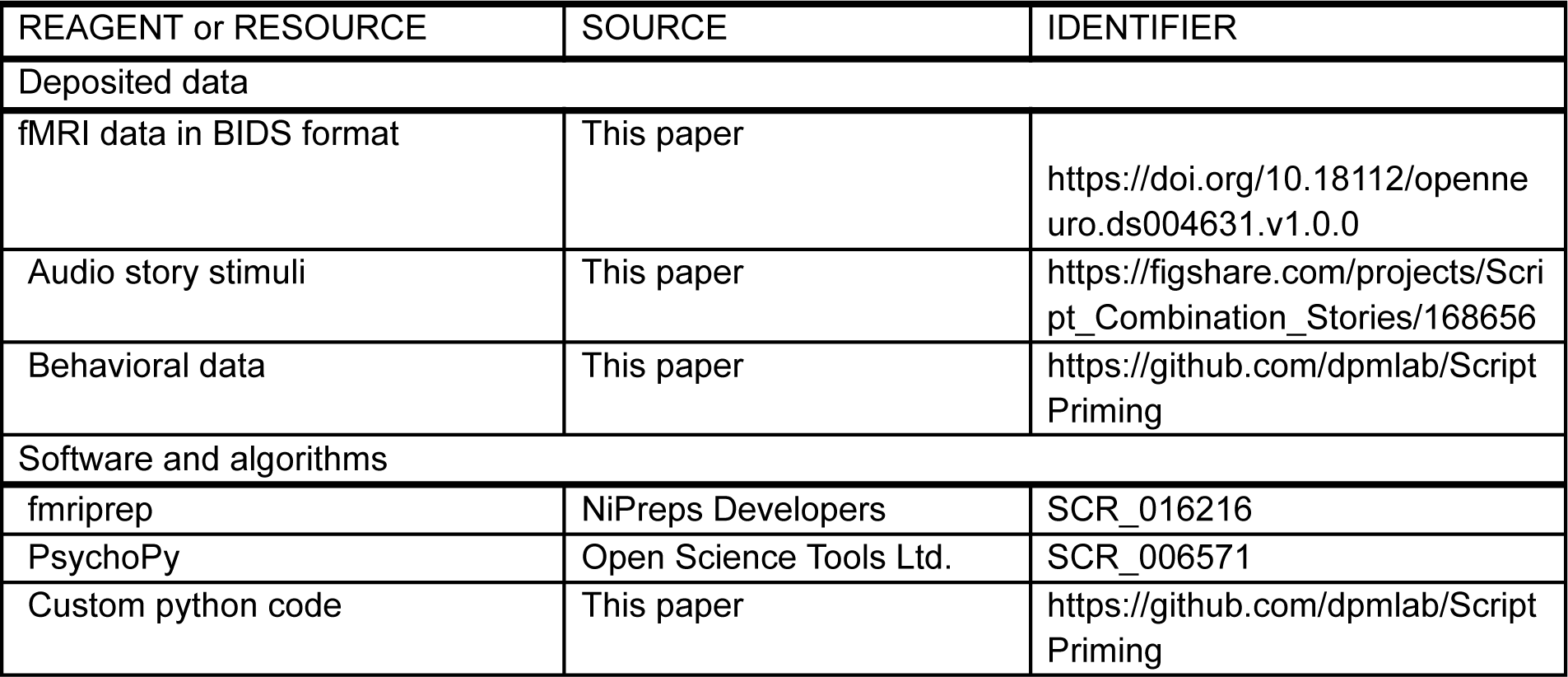

